## Supporting Information for "Analysis of Conformational Exchange Processes using Methyl-TROSY-Based Hahn Echo Measurements of Quadruple-Quantum Relaxation"

### Table of Contents

*Supporting Text*

Analysis of multi-state exchange processes

*Supporting Figures*

Fig. S1 Assigned 2D spectrum of FLN5 acquired using QQ Hahn echo experiment

Fig. S2 Pseudo-3D lineshape fitting for the measurement of $^{13}$C CSA and $S^2_{axis}\tau_c$

Fig. S3 Comparison of measurements of methyl $S^2_{axis}\tau_c$ parameters

Fig. S4 Constraints on $(\xi_H, \xi_C)$ parameter space arising from HE measurements

Fig. S5 Illustration of the effect of three-state chemical exchange on the analysis of multiple-quantum HE measurements

Fig. S6 Global fitting of HE and CPMG measurements

*Supporting Tables*

Table S1 Measured methyl DQ' relaxation rates for FLN5, 283 K, 600 to 950 MHz

Table S2 Measured methyl QQ relaxation rates for FLN5, 283 K, 600 to 950 MHz

Table S3 Measured methyl ZQ relaxation rates for FLN5, 283 K, 600 to 950 MHz

Table S4 Measured methyl DQ relaxation rates for FLN5, 283 K, 600 to 950 MHz

Table S5 Measured methyl $^{13}$C CSA, $^1$H CSA and $S^2_{axis}\tau_c$ values for FLN5, 283 K

Table S6 Fitted chemical shift perturbations for FLN5 excited states, 283 K

*Supporting Listings*

Listing S1 Bruker format pulse sequence for measurement of methyl Hahn echo DQ' and QQ relaxation

Listing S2 Processing scripts for analysis of DQ' and QQ Hahn echo experiments

Listing S3 Bruker format pulse sequence for measurement of methyl $^{13}$C CSA and $S^2_{axis}\tau_c$

Listing S4 Bruker format pulse sequence for measurement of methyl $^1$H CSA

*References*

### Supporting Text

*Analysis of multi-state exchange processes*

In the presence of multiple, uncorrelated chemical exchange processes, the exchange contribution to relaxation will contain terms arising from each process (here assuming fast chemical exchange):

$$R_{ex} = \sum_i \left(\xi_C^{(i)} + n\xi_H^{(i)}\right)^2 B_0^2 \quad (1)$$

where $\xi_C^{(i)}$ and $\xi_H^{(i)}$ (Eq. 3, main text) represent the *i*-th exchange process, and $n$ varies depending on the multiple quantum coherence being considered.

Given HE measurements of a single coherence, the presence of multiple exchange processes cannot be distinguished from a single process, as the functional form of the observed relaxation rate, $R_{2,obs} = R_{2,0} + \beta B_0^2$, is identical. Moreover, the situation is not improved if two measurements (e.g. ZQ and DQ) are made, as apparent two-state parameters $\xi_H^{app}$ and $\xi_C^{app}$ can always be determined that are consistent with observations:

$$\begin{aligned} \beta_{ZQ} &= (\xi_C^1 - \xi_H^1)^2 + (\xi_C^2 - \xi_H^2)^2 = \left(\xi_C^{app} - \xi_H^{app}\right)^2 \\ \beta_{DQ} &= (\xi_C^1 + \xi_H^1)^2 + (\xi_C^2 + \xi_H^2)^2 = \left(\xi_C^{app} + \xi_H^{app}\right)^2 \end{aligned} \quad (2)$$

However, if additional measurements (e.g. DQ' and QQ) are also available, in general a consistent apparent two-state solution will not exist:

$$\begin{aligned} \beta_{ZQ} &= (\xi_C^1 - \xi_H^1)^2 + (\xi_C^2 - \xi_H^2)^2 \neq \left(\xi_C^{app} - \xi_H^{app}\right)^2 \\ \beta_{DQ} &= (\xi_C^1 + \xi_H^1)^2 + (\xi_C^2 + \xi_H^2)^2 \neq \left(\xi_C^{app} + \xi_H^{app}\right)^2 \\ \beta_{DQ'} &= (\xi_C^1 - 3\xi_H^1)^2 + (\xi_C^2 - 3\xi_H^2)^2 \neq \left(\xi_C^{app} - 3\xi_H^{app}\right)^2 \\ \beta_{QQ} &= (\xi_C^1 + 3\xi_H^1)^2 + (\xi_C^2 + 3\xi_H^2)^2 \neq \left(\xi_C^{app} + 3\xi_H^{app}\right)^2 \end{aligned} \quad (3)$$

At a graphical level, this corresponds to the non-intersection of constraints on $(\xi_H, \xi_C)$ parameter space, illustrated in Fig. S5. We suggest that this analysis may serve as a useful tool for the detection of such occurrences.

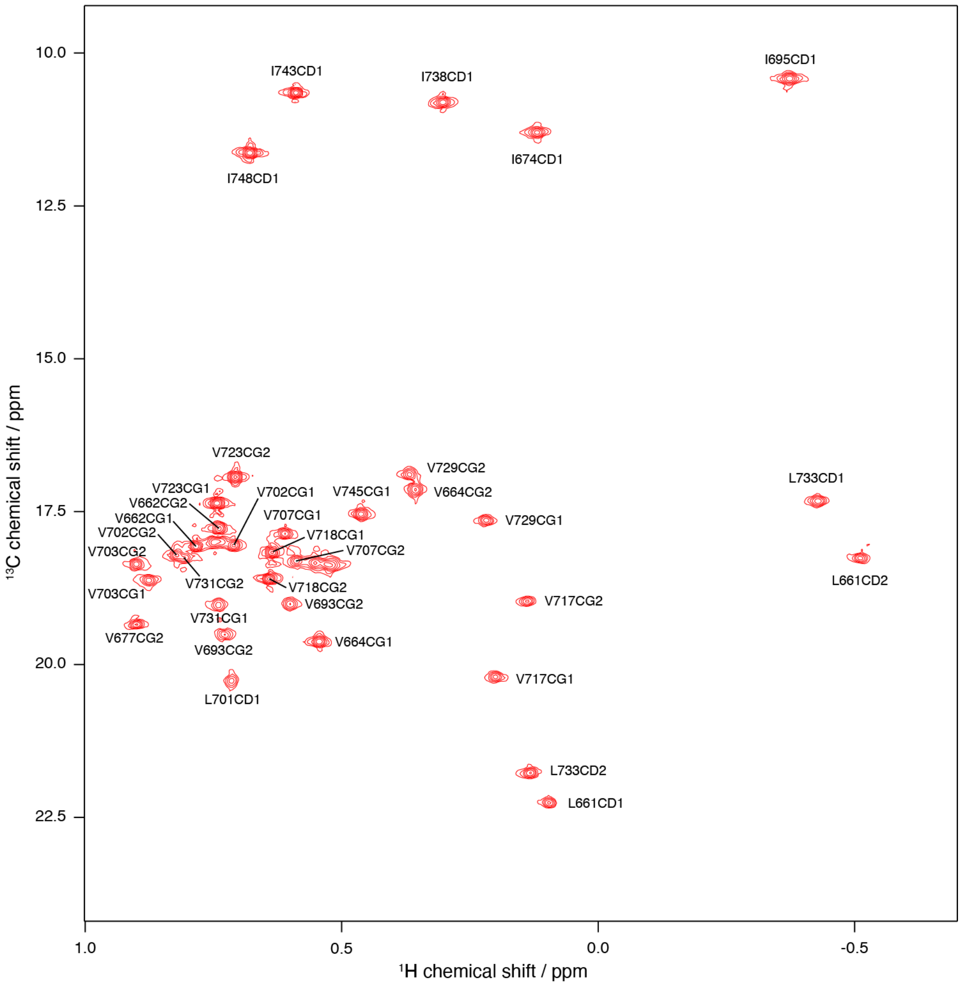


**Figure S1.** Assigned 2D $^{1}H$,$^{13}C$ correlation spectrum of FLN5 acquired using QQ Hahn echo experiment (Fig. 2a) with a 0.1 ms relaxation delay.

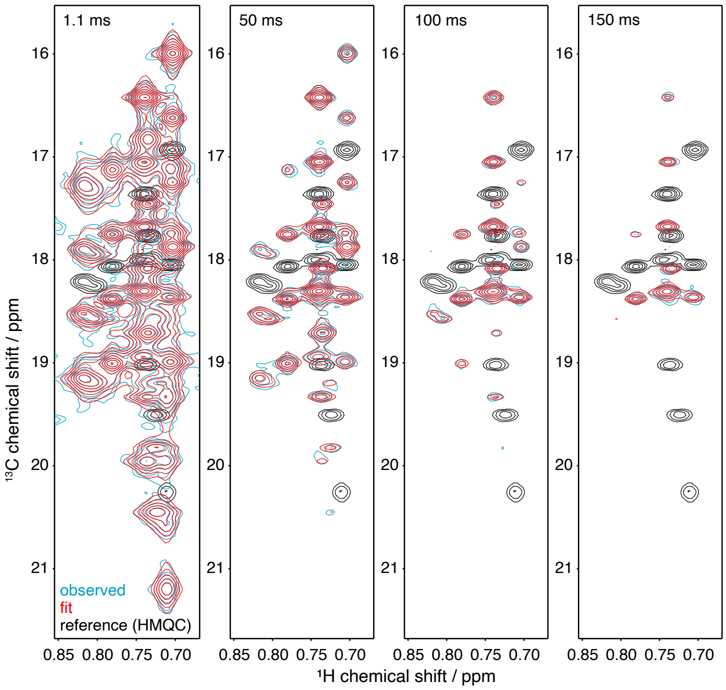


**Figure S2.** Pseudo-3D lineshape fitting of spectra obtained using the pulse sequence in Fig. 3a, for the measurement of $^{13}$C CSA and $S^2_{axis}\tau_c$ values in FLN5, 800 MHz, 283 K. The fitting of a cluster of overlapped resonances (red) to the observed spectrum (blue) is shown, with relaxation times as indicated (top). For reference, an HMQC spectrum is also plotted (black).

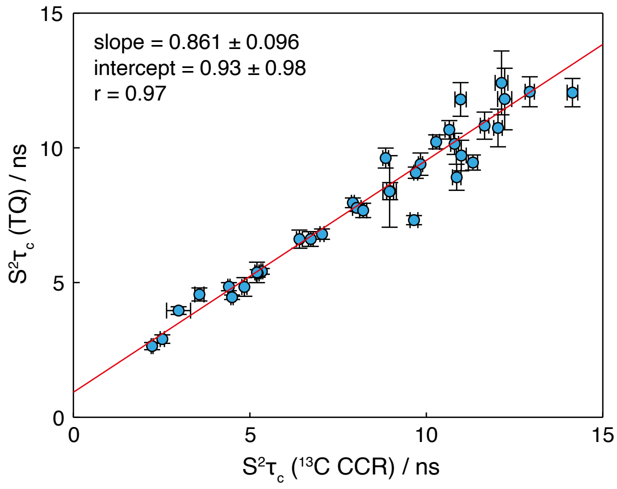


**Figure S3.** Comparison of measurements of methyl $S^2_{axis}\tau_c$ values in FLN5, 283 K, via the 2D lineshape fitting of $^{13}$C multiplets (Fig. 3A), and via $^{1}$H TQ build-up experiments[1]. Error bars indicate the standard error derived from fitting.

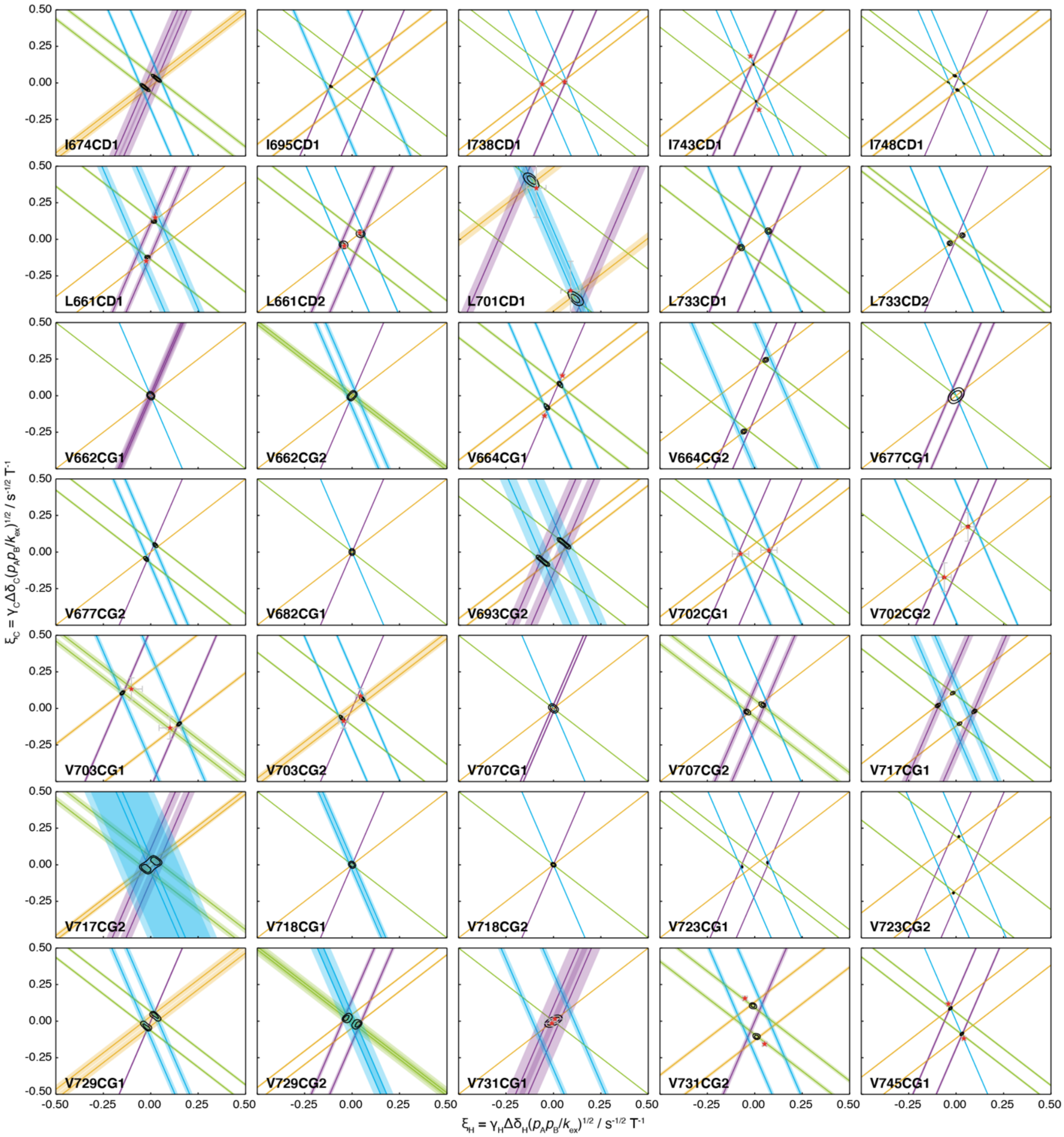


**Figure S4.** Constraints on $(\xi_H, \xi_C)$ parameter space arising from HE measurements, plotted for all methyl resonances. Straight lines indicate values of $\xi_H$ and $\xi_C$ obtained from linear regression of HE measurements, calculated according to Table 1 assuming fast exchange and subtracting measured CSA contributions. Shading indicates the standard error propagated from linear regression analysis and CSA measurements. Black contours indicate 68 and 95% confidence intervals in $\xi_H$ and $\xi_C$, based on all four HE measurements and assuming two-state fast exchange. Red symbols indicate $\xi_H$ and $\xi_C$ parameters derived from global fitting of HE and CPMG data (Fig. S5).

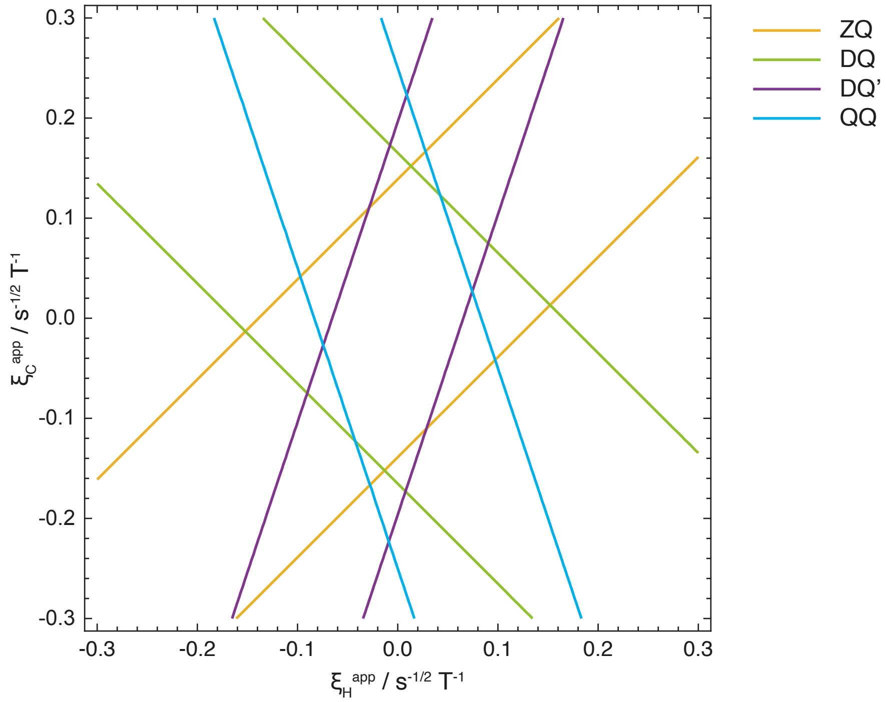


**Figure S5.** Illustration of the effect of three-state chemical exchange on the analysis of multiple-quantum HE measurements. The non-intersecting constraints that arise on two-state $(\xi_H, \xi_C)$ parameter space are illustrated for two non-correlated exchange processes, with $\xi_H^{(1)}$ = 0.05 s$^{-1/2}$ T$^{-1}$, $\xi_C^{(1)}$ = 0.1 s$^{-1/2}$ T$^{-1}$, $\xi_H^{(2)}$ = 0.03 s$^{-1/2}$ T$^{-1}$ and $\xi_C^{(2)}$ = -0.1 s$^{-1/2}$ T$^{-1}$.

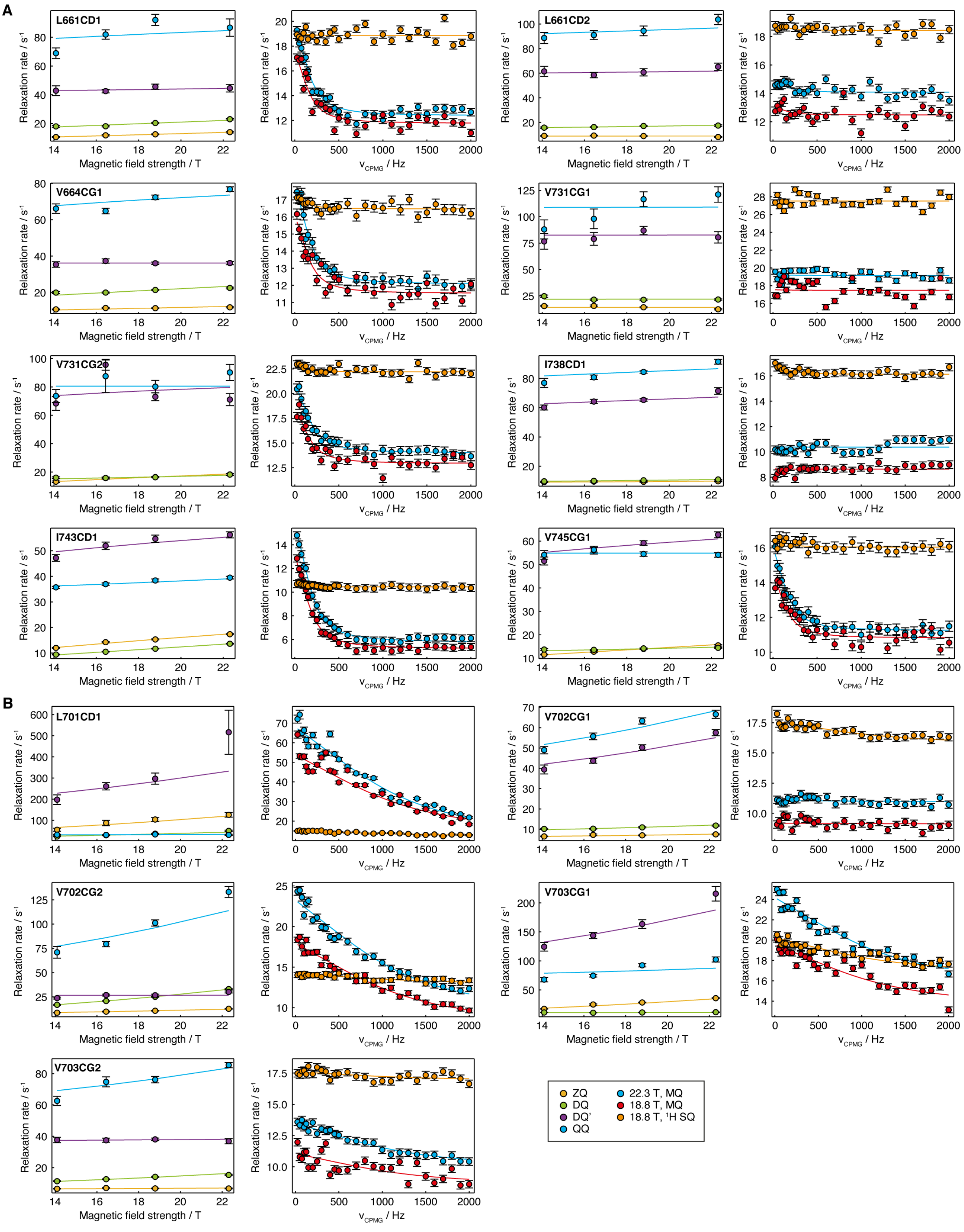


**Figure S6.** Global fitting of HE and CPMG measurements. Panels (A) and (B) show methyl groups in each of the two exchange clusters identified and shown in Fig. 6.

| methyl | 14.1 T | 16.4 T | 18.8 T | 22.3 T |
|---|---|---|---|---|
| I674CD1 | 64.9 ± 3.0 | 69.7 ± 1.8 | 71.3 ± 1.8 | 68.9 ± 2.1 |
| I695CD1 | 81.5 ± 1.8 | 89.7 ± 1.6 | 97.0 ± 1.4 | 115.7 ± 3.6 |
| I738CD1 | 60.2 ± 1.7 | 64.3 ± 1.5 | 65.41 ± 0.93 | 71.6 ± 2.2 |
| I743CD1 | 47.2 ± 1.3 | 52.0 ± 1.4 | 54.7 ± 1.4 | 56.3 ± 1.2 |
| I748CD1 | 25.8 ± 1.5 | 26.5 ± 1.4 | 26.7 ± 1.5 | 25.6 ± 1.6 |
| L661CD1 | 43.1 ± 3.4 | 42.9 ± 1.2 | 46.1 ± 1.8 | 45.3 ± 2.6 |
| L661CD2 | 62.1 ± 4.1 | 59.4 ± 2.2 | 61.9 ± 3.0 | 66.8 ± 3.0 |
| L701CD1 | 198.0 ± 23.0 | 261.0 ± 17.0 | 297.0 ± 27.0 | 520.0 ± 100.0 |
| L733CD1 | 49.3 ± 2.6 | 53.1 ± 1.8 | 59.9 ± 1.9 | 63.8 ± 2.7 |
| L733CD2 | 33.0 ± 1.4 | 34.5 ± 1.1 | 34.88 ± 0.97 | 33.4 ± 1.0 |
| V662CG1 | 22.5 ± 0.7 | 22.02 ± 0.76 | 22.81 ± 0.66 | 22.69 ± 0.77 |
| V662CG2 | 25.0 ± 1.1 | 25.95 ± 0.92 | 25.07 ± 0.84 | 24.37 ± 0.86 |
| V664CG1 | 36.1 ± 1.5 | 38.3 ± 1.2 | 37.34 ± 0.67 | 38.0 ± 0.9 |
| V664CG2 | 44.4 ± 1.8 | 46.0 ± 1.0 | 49.04 ± 0.93 | 51.6 ± 1.4 |
| V677CG1 | 45.6 ± 3.5 | 47.3 ± 2.4 | 46.4 ± 2.0 | 49.4 ± 2.3 |
| V677CG2 | 35.3 ± 1.4 | 34.3 ± 1.5 | 34.2 ± 1.2 | 32.6 ± 1.0 |
| V682CG1 | 29.9 ± 1.1 | 24.7 ± 1.6 | 29.58 ± 0.97 | 25.8 ± 1.3 |
| V693CG2 | 71.1 ± 4.4 | 81.5 ± 3.9 | 75.8 ± 2.3 | 83.7 ± 6.2 |
| V702CG1 | 39.8 ± 2.2 | 44.2 ± 1.1 | 50.9 ± 1.4 | 58.5 ± 1.5 |
| V702CG2 | 24.2 ± 0.9 | 27.57 ± 0.99 | 27.6 ± 1.1 | 31.3 ± 1.0 |
| V703CG1 | 124.9 ± 7.6 | 144.5 ± 5.1 | 164.4 ± 7.4 | 217.0 ± 13.0 |
| V703CG2 | 38.3 ± 1.7 | 38.3 ± 1.3 | 39.14 ± 0.78 | 38.4 ± 1.7 |
| V707CG1 | 41.3 ± 1.5 | 41.66 ± 0.81 | 41.88 ± 0.64 | 42.1 ± 1.3 |
| V707CG2 | 50.4 ± 2.1 | 47.6 ± 1.7 | 49.1 ± 1.2 | 54.6 ± 1.5 |
| V717CG1 | 61.7 ± 5.0 | 73.9 ± 5.9 | 85.0 ± 2.0 | 87.0 ± 5.2 |
| V717CG2 | 64.6 ± 5.2 | 69.0 ± 4.7 | 73.3 ± 1.6 | 71.5 ± 3.3 |
| V718CG1 | 24.8 ± 0.8 | 25.26 ± 0.65 | 25.37 ± 0.72 | 24.64 ± 0.8 |
| V718CG2 | 22.2 ± 0.8 | 22.79 ± 0.75 | 23.13 ± 0.75 | 22.37 ± 0.75 |
| V723CG1 | 33.8 ± 0.9 | 36.78 ± 0.89 | 40.72 ± 0.99 | 46.88 ± 0.86 |
| V723CG2 | 38.1 ± 1.3 | 43.2 ± 1.1 | 48.4 ± 1.0 | 59.4 ± 1.0 |
| V729CG1 | 53.6 ± 3.1 | 59.7 ± 1.5 | 56.7 ± 1.7 | 57.7 ± 2.5 |
| V729CG2 | 56.3 ± 1.5 | 60.2 ± 2.6 | 59.06 ± 0.85 | 65.3 ± 2.3 |
| V731CG1 | 78.1 ± 7.4 | 81.0 ± 6.1 | 89.6 ± 3.9 | 84.2 ± 5.3 |
| V731CG2 | 69.1 ± 4.9 | 96.6 ± 3.9 | 74.4 ± 2.7 | 72.9 ± 4.2 |
| V745CG1 | 51.9 ± 1.7 | 56.6 ± 1.6 | 59.82 ± 0.94 | 63.6 ± 1.2 |

**Table S1.** Measured methyl DQ’ relaxation rates for FLN5, 283 K, 600 to 950 MHz.

| **methyl** | **14.1 T** | **16.4 T** | **18.8 T** | **22.3 T** |
|---|---|---|---|---|
| I674CD1 | 91.6 ± 4.8 | 91.1 ± 3.2 | 93.2 ± 3.0 | 100.2 ± 3.6 |
| I695CD1 | 115.2 ± 4.2 | 121.4 ± 3.0 | 143.1 ± 1.6 | 166.5 ± 6.3 |
| I738CD1 | 77.6 ± 3.1 | 81.7 ± 1.6 | 85.63 ± 0.84 | 93.1 ± 1.6 |
| I743CD1 | 35.9 ± 0.4 | 37.23 ± 0.49 | 38.75 ± 0.58 | 40.02 ± 0.5 |
| I748CD1 | 27.5 ± 0.6 | 28.42 ± 0.81 | 29.53 ± 0.63 | 31.6 ± 1.1 |
| L661CD1 | 71.2 ± 3.7 | 85.0 ± 3.2 | 96.1 ± 3.9 | 92.5 ± 6.0 |
| L661CD2 | 89.9 ± 4.5 | 92.7 ± 3.6 | 96.6 ± 4.0 | 106.8 ± 4.2 |
| L701CD1 | 33.4 ± 1.9 | 32.9 ± 1.4 | 35.99 ± 0.96 | 34.8 ± 2.3 |
| L733CD1 | 90.2 ± 5.1 | 90.7 ± 4.4 | 96.4 ± 3.8 | 109.5 ± 5.1 |
| L733CD2 | 51.4 ± 2.6 | 57.54 ± 0.88 | 60.8 ± 1.6 | 72.6 ± 1.7 |
| V662CG1 | 29.7 ± 0.9 | 29.74 ± 0.6 | 31.49 ± 0.56 | 30.43 ± 0.69 |
| V662CG2 | 34.9 ± 1.6 | 33.1 ± 1.1 | 36.1 ± 0.45 | 37.3 ± 0.84 |
| V664CG1 | 66.3 ± 2.6 | 65.0 ± 1.5 | 72.7 ± 1.2 | 77.3 ± 1.3 |
| V664CG2 | 121.9 ± 8.0 | 131.8 ± 3.3 | 144.4 ± 2.9 | 194.1 ± 4.6 |
| V677CG1 | 67.3 ± 4.7 | 52.6 ± 2.6 | 57.1 ± 1.9 | 54.3 ± 3.0 |
| V677CG2 | 56.1 ± 1.8 | 55.1 ± 1.4 | 56.1 ± 1.3 | 59.4 ± 1.8 |
| V682CG1 | 42.2 ± 1.4 | 37.1 ± 3.0 | 40.5 ± 1.4 | 36.1 ± 1.2 |
| V693CG2 | 99.6 ± 7.5 | 103.4 ± 6.4 | 122.8 ± 4.0 | 114.2 ± 5.7 |
| V702CG1 | 50.3 ± 1.8 | 57.5 ± 1.7 | 65.7 ± 1.5 | 69.9 ± 2.0 |
| V702CG2 | 72.3 ± 6.1 | 81.4 ± 2.7 | 103.7 ± 3.2 | 136.7 ± 5.7 |
| V703CG1 | 68.9 ± 3.8 | 76.0 ± 2.3 | 94.4 ± 2.7 | 105.2 ± 4.4 |
| V703CG2 | 64.8 ± 2.9 | 77.8 ± 3.2 | 80.1 ± 2.0 | 91.0 ± 1.6 |
| V707CG1 | 63.7 ± 2.0 | 56.8 ± 2.5 | 60.0 ± 1.3 | 61.5 ± 2.9 |
| V707CG2 | 67.6 ± 3.2 | 71.5 ± 2.3 | 78.1 ± 2.7 | 84.1 ± 4.2 |
| V717CG1 | 88.4 ± 7.6 | 84.9 ± 4.6 | 84.5 ± 2.4 | 93.3 ± 3.9 |
| V717CG2 | 95.2 ± 8.8 | 93.1 ± 4.0 | 107.0 ± 4.7 | 92.2 ± 6.2 |
| V718CG1 | 35.7 ± 0.7 | 37.07 ± 0.9 | 36.68 ± 0.51 | 36.93 ± 0.83 |
| V718CG2 | 30.4 ± 0.6 | 31.58 ± 0.63 | 31.57 ± 0.57 | 30.42 ± 0.76 |
| V723CG1 | 50.2 ± 0.9 | 53.27 ± 0.74 | 58.78 ± 0.77 | 67.2 ± 1.2 |
| V723CG2 | 52.4 ± 1.1 | 57.96 ± 0.96 | 64.6 ± 0.89 | 78.5 ± 1.2 |
| V729CG1 | 82.0 ± 5.0 | 81.5 ± 2.4 | 87.1 ± 3.5 | 86.0 ± 3.1 |
| V729CG2 | 64.8 ± 4.4 | 68.9 ± 2.9 | 72.6 ± 4.4 | 69.6 ± 2.3 |
| V731CG1 | 89.2 ± 9.0 | 99.6 ± 9.3 | 118.7 ± 7.0 | 124.1 ± 7.2 |
| V731CG2 | 75.0 ± 4.5 | 89.0 ± 11.0 | 82.6 ± 4.4 | 93.6 ± 5.7 |
| V745CG1 | 54.7 ± 1.8 | 57.1 ± 1.0 | 55.52 ± 0.95 | 55.55 ± 0.97 |

**Table S2.** Measured methyl QQ relaxation rates for FLN5, 283 K, 600 to 950 MHz.

| methyl | 14.1 T | 16.4 T | 18.8 T | 22.3 T |
|---|---|---|---|---|
| I674CD1 | 10.11 ± 0.2 | 10.56 ± 0.25 | 10.32 ± 0.23 | 10.56 ± 0.28 |
| I695CD1 | 9.63 ± 0.16 | 10.69 ± 0.25 | 10.71 ± 0.28 | 11.75 ± 0.34 |
| I738CD1 | 9.38 ± 0.23 | 9.81 ± 0.41 | 9.72 ± 0.31 | 10.21 ± 0.33 |
| I743CD1 | 12.07 ± 0.16 | 14.33 ± 0.21 | 15.44 ± 0.23 | 17.52 ± 0.22 |
| I748CD1 | 4.652 ± 0.097 | 5.81 ± 0.38 | 5.25 ± 0.19 | 5.89 ± 0.22 |
| L661CD1 | 11.35 ± 0.24 | 12.84 ± 0.48 | 13.87 ± 0.31 | 15.94 ± 0.49 |
| L661CD2 | 9.83 ± 0.24 | 10.33 ± 0.31 | 10.3 ± 0.3 | 10.07 ± 0.32 |
| L701CD1 | 55.6 ± 7.3 | 88.0 ± 13.0 | 104.5 ± 9.1 | 127.0 ± 11.0 |
| L733CD1 | 7.8 ± 0.3 | 8.13 ± 0.3 | 7.88 ± 0.24 | 7.65 ± 0.27 |
| L733CD2 | 8.63 ± 0.24 | 8.39 ± 0.33 | 8.04 ± 0.2 | 8.52 ± 0.22 |
| V662CG1 | 5.51 ± 0.14 | 6.19 ± 0.28 | 5.85 ± 0.15 | 6.12 ± 0.16 |
| V662CG2 | 6.14 ± 0.19 | 7.24 ± 0.36 | 6.79 ± 0.18 | 7.15 ± 0.18 |
| V664CG1 | 11.19 ± 0.28 | 12.15 ± 0.31 | 12.27 ± 0.21 | 13.13 ± 0.23 |
| V664CG2 | 18.21 ± 0.83 | 21.56 ± 0.52 | 24.84 ± 0.4 | 30.1 ± 0.66 |
| V677CG1 | 16.18 ± 0.52 | 16.3 ± 0.47 | 15.74 ± 0.54 | 15.63 ± 0.4 |
| V677CG2 | 6.94 ± 0.23 | 7.66 ± 0.32 | 7.55 ± 0.15 | 7.69 ± 0.21 |
| V682CG1 | 7.7 ± 0.19 | 8.46 ± 0.28 | 7.47 ± 0.12 | 7.92 ± 0.25 |
| V693CG2 | 12.45 ± 0.43 | 13.04 ± 0.21 | 13.56 ± 0.38 | 13.95 ± 0.29 |
| V702CG1 | 7.08 ± 0.18 | 8.07 ± 0.42 | 8.09 ± 0.23 | 9.03 ± 0.25 |
| V702CG2 | 9.38 ± 0.21 | 10.79 ± 0.45 | 12.01 ± 0.27 | 14.55 ± 0.27 |
| V703CG1 | 17.93 ± 0.36 | 25.7 ± 1.0 | 29.5 ± 0.48 | 37.38 ± 0.94 |
| V703CG2 | 7.57 ± 0.18 | 8.42 ± 0.33 | 8.6 ± 0.23 | 9.34 ± 0.22 |
| V707CG1 | 8.63 ± 0.29 | 8.68 ± 0.42 | 8.94 ± 0.27 | 9.49 ± 0.23 |
| V707CG2 | 8.32 ± 0.21 | 9.12 ± 0.49 | 8.82 ± 0.22 | 8.64 ± 0.21 |
| V717CG1 | 14.32 ± 0.41 | 16.22 ± 0.26 | 17.33 ± 0.33 | 19.87 ± 0.34 |
| V717CG2 | 11.51 ± 0.29 | 11.96 ± 0.32 | 12.14 ± 0.27 | 12.59 ± 0.2 |
| V718CG1 | 5.899 ± 0.089 | 6.61 ± 0.28 | 6.38 ± 0.14 | 6.39 ± 0.21 |
| V718CG2 | 5.59 ± 0.1 | 6.09 ± 0.37 | 5.8 ± 0.13 | 5.75 ± 0.14 |
| V723CG1 | 7.3 ± 0.18 | 7.62 ± 0.33 | 7.32 ± 0.2 | 7.48 ± 0.14 |
| V723CG2 | 13.27 ± 0.27 | 16.19 ± 0.33 | 18.8 ± 0.45 | 22.82 ± 0.34 |
| V729CG1 | 9.84 ± 0.24 | 11.03 ± 0.29 | 10.52 ± 0.24 | 11.18 ± 0.23 |
| V729CG2 | 9.2 ± 0.36 | 9.66 ± 0.37 | 9.99 ± 0.26 | 11.08 ± 0.31 |
| V731CG1 | 16.79 ± 0.67 | 17.46 ± 0.79 | 16.75 ± 0.37 | 15.84 ± 0.37 |
| V731CG2 | 14.13 ± 0.53 | 16.99 ± 0.73 | 18.04 ± 0.37 | 20.6 ± 0.75 |
| V745CG1 | 12.02 ± 0.16 | 13.39 ± 0.27 | 14.75 ± 0.19 | 16.57 ± 0.29 |

**Table S3.** Measured methyl ZQ relaxation rates for FLN5, 283 K, 600 to 950 MHz.

| methyl | 14.1 T | 16.4 T | 18.8 T | 22.3 T |
|---|---|---|---|---|
| I674CD1 | 12.84 ± 0.2 | 12.702 ± 0.068 | 13.76 ± 0.2 | 13.9 ± 0.16 |
| I695CD1 | 13.64 ± 0.23 | 14.905 ± 0.083 | 16.91 ± 0.17 | 19.89 ± 0.22 |
| I738CD1 | 10.12 ± 0.12 | 10.576 ± 0.068 | 11.11 ± 0.13 | 11.98 ± 0.1 |
| I743CD1 | 9.53 ± 0.15 | 10.658 ± 0.076 | 11.91 ± 0.12 | 13.94 ± 0.14 |
| I748CD1 | 4.206 ± 0.095 | 4.313 ± 0.078 | 4.494 ± 0.071 | 5.063 ± 0.058 |
| L661CD1 | 19.44 ± 0.64 | 20.06 ± 0.44 | 22.92 ± 0.33 | 26.61 ± 0.45 |
| L661CD2 | 16.7 ± 0.55 | 17.5 ± 0.28 | 18.84 ± 0.44 | 19.88 ± 0.58 |
| L701CD1 | 24.4 ± 1.4 | 30.4 ± 1.8 | 38.1 ± 2.5 | 52.2 ± 2.9 |
| L733CD1 | 18.54 ± 0.69 | 19.73 ± 0.34 | 22.23 ± 0.53 | 24.82 ± 0.79 |
| L733CD2 | 13.09 ± 0.3 | 13.41 ± 0.24 | 14.28 ± 0.25 | 15.46 ± 0.26 |
| V662CG1 | 7.58 ± 0.18 | 7.67 ± 0.14 | 8.1 ± 0.14 | 8.21 ± 0.13 |
| V662CG2 | 8.97 ± 0.22 | 9.21 ± 0.17 | 9.53 ± 0.18 | 10.43 ± 0.16 |
| V664CG1 | 20.26 ± 0.58 | 20.41 ± 0.3 | 21.97 ± 0.3 | 23.33 ± 0.47 |
| V664CG2 | 34.8 ± 1.5 | 39.3 ± 1.3 | 46.67 ± 0.76 | 57.4 ± 2.1 |
| V677CG1 | 33.2 ± 1.6 | 34.3 ± 1.3 | 33.8 ± 1.0 | 34.7 ± 1.1 |
| V677CG2 | 11.38 ± 0.26 | 12.1 ± 0.15 | 13.2 ± 0.16 | 14.07 ± 0.21 |
| V682CG1 | 10.76 ± 0.25 | 10.5 ± 0.19 | 10.59 ± 0.16 | 11.12 ± 0.12 |
| V693CG2 | 17.86 ± 0.27 | 18.72 ± 0.25 | 20.36 ± 0.27 | 22.86 ± 0.52 |
| V702CG1 | 11.08 ± 0.28 | 11.5 ± 0.19 | 12.57 ± 0.25 | 14.15 ± 0.28 |
| V702CG2 | 17.84 ± 0.56 | 22.18 ± 0.31 | 26.89 ± 0.44 | 35.67 ± 0.55 |
| V703CG1 | 11.96 ± 0.2 | 11.75 ± 0.24 | 12.66 ± 0.21 | 13.75 ± 0.36 |
| V703CG2 | 12.92 ± 0.29 | 14.76 ± 0.2 | 16.95 ± 0.36 | 19.29 ± 0.36 |
| V707CG1 | 12.47 ± 0.34 | 12.37 ± 0.23 | 13.56 ± 0.24 | 13.45 ± 0.17 |
| V707CG2 | 13.28 ± 0.38 | 13.08 ± 0.12 | 14.2 ± 0.2 | 14.7 ± 0.29 |
| V717CG1 | 17.91 ± 0.31 | 18.38 ± 0.34 | 19.38 ± 0.27 | 21.56 ± 0.4 |
| V717CG2 | 17.12 ± 0.33 | 16.7 ± 0.18 | 17.8 ± 0.42 | 18.62 ± 0.43 |
| V718CG1 | 8.89 ± 0.22 | 9.22 ± 0.14 | 9.54 ± 0.2 | 9.24 ± 0.13 |
| V718CG2 | 7.97 ± 0.21 | 7.79 ± 0.1 | 8.12 ± 0.16 | 7.64 ± 0.12 |
| V723CG1 | 11.5 ± 0.24 | 11.69 ± 0.17 | 11.93 ± 0.19 | 12.64 ± 0.2 |
| V723CG2 | 16.33 ± 0.45 | 19.62 ± 0.28 | 22.46 ± 0.2 | 28.73 ± 0.54 |
| V729CG1 | 14.25 ± 0.37 | 14.84 ± 0.14 | 15.88 ± 0.26 | 16.49 ± 0.28 |
| V729CG2 | 11.59 ± 0.23 | 11.76 ± 0.13 | 12.47 ± 0.23 | 12.99 ± 0.2 |
| V731CG1 | 26.1 ± 1.0 | 23.25 ± 0.57 | 23.65 ± 0.72 | 24.87 ± 0.45 |
| V731CG2 | 17.13 ± 0.58 | 17.31 ± 0.28 | 18.29 ± 0.19 | 20.99 ± 0.44 |
| V745CG1 | 14.24 ± 0.41 | 14.27 ± 0.12 | 15.1 ± 0.22 | 15.7 ± 0.19 |

**Table S4.** Measured methyl DQ relaxation rates for FLN5, 283 K, 600 to 950 MHz.

| methyl | $S^2\tau_c$ ([13]C CCR) / ns | $S^2\tau_c$ (TQ) / ns | [13]C CSA / ppm | [1]H CSA / ppm |
|---|---|---|---|---|
| I674CD1 | 9.83 ± 0.06 | 9.39 ± 0.42 | 15.78 ± 0.30 | 0.25 ± 0.10 |
| I695CD1 | 7.92 ± 0.04 | 7.96 ± 0.18 | 24.15 ± 0.29 | 1.55 ± 0.10 |
| I738CD1 | 9.70 ± 0.06 | 9.08 ± 0.21 | 19.75 ± 0.29 | 0.60 ± 0.06 |
| I743CD1 | 4.50 ± 0.03 | 4.46 ± 0.08 | 16.34 ± 0.39 | 0.22 ± 0.05 |
| I748CD1 | 5.34 ± 0.03 | 5.42 ± 0.11 | 20.00 ± 0.26 | 0.45 ± 0.07 |
| L661CD1 | 10.97 ± 0.16 | 11.80 ± 0.63 | 33.85 ± 0.95 | 1.14 ± 0.10 |
| L661CD2 | 8.97 ± 0.10 | 8.39 ± 0.32 | 36.26 ± 0.66 | 0.31 ± 0.12 |
| L701CD1 | 2.98 ± 0.34 | 3.96 ± 0.14 | 44.60 ± 8.10 | 0.61 ± 0.15 |
| L733CD1 | 6.40 ± 0.08 | 6.61 ± 0.34 | 41.28 ± 0.78 | 0.88 ± 0.15 |
| L733CD2 | 6.73 ± 0.07 | 6.62 ± 0.27 | 43.60 ± 0.59 | 1.02 ± 0.10 |
| V662CG1 | 7.04 ± 0.06 | 6.80 ± 0.19 | 29.49 ± 0.45 | 0.37 ± 0.06 |
| V662CG2 | 8.85 ± 0.08 | 9.62 ± 0.37 | 33.45 ± 0.57 | 0.36 ± 0.05 |
| V664CG1 | 4.84 ± 0.07 | 4.84 ± 0.35 | 33.10 ± 1.00 | -0.30 ± 0.06 |
| V664CG2 | 3.57 ± 0.13 | 4.56 ± 0.24 | 44.80 ± 2.20 | 0.21 ± 0.13 |
| V677CG1 | 8.96 ± 0.19 | 8.38 ± 1.33 | 28.50 ± 1.20 | 0.78 ± 0.22 |
| V677CG2 | 10.80 ± 0.11 | 10.15 ± 0.39 | 26.86 ± 0.59 | 0.25 ± 0.09 |
| V682CG1 | 9.65 ± 0.11 | 7.32 ± 0.17 | 28.82 ± 0.63 | 0.38 ± 0.10 |
| V693CG2 | 10.99 ± 0.14 | 9.71 ± 0.57 | 26.30 ± 0.79 | 0.14 ± 0.17 |
| V702CG1 | 10.28 ± 0.10 | 10.22 ± 0.26 | 30.81 ± 0.54 | 0.58 ± 0.08 |
| V702CG2 | 8.03 ± 0.12 | 7.77 ± 0.13 | 38.28 ± 0.89 | 0.43 ± 0.04 |
| V703CG1 | 8.21 ± 0.10 | 7.68 ± 0.27 | 35.19 ± 0.81 | 0.36 ± 0.19 |
| V703CG2 | 11.65 ± 0.12 | 10.82 ± 0.50 | 38.90 ± 0.66 | 0.78 ± 0.07 |
| V707CG1 | 10.65 ± 0.12 | 10.67 ± 0.34 | 32.97 ± 0.72 | 0.78 ± 0.09 |
| V707CG2 | 10.86 ± 0.12 | 8.91 ± 0.48 | 29.32 ± 0.62 | 0.49 ± 0.11 |
| V717CG1 | 11.32 ± 0.13 | 9.45 ± 0.28 | 29.60 ± 0.73 | 0.36 ± 0.10 |
| V717CG2 | 14.14 ± 0.15 | 12.05 ± 0.53 | 25.54 ± 0.62 | 0.03 ± 0.22 |
| V718CG1 | 5.22 ± 0.04 | 5.34 ± 0.14 | 32.65 ± 0.50 | 0.04 ± 0.06 |
| V718CG2 | 4.41 ± 0.03 | 4.85 ± 0.15 | 33.72 ± 0.44 | 0.14 ± 0.05 |
| V723CG1 | 2.23 ± 0.03 | 2.64 ± 0.13 | 27.94 ± 0.65 | -0.04 ± 0.05 |
| V723CG2 | 2.52 ± 0.06 | 2.90 ± 0.16 | 44.80 ± 1.30 | -0.12 ± 0.02 |
| V729CG1 | 12.03 ± 0.12 | 10.74 ± 0.70 | 26.60 ± 0.61 | 0.41 ± 0.14 |
| V729CG2 | 12.93 ± 0.13 | 12.08 ± 0.56 | 30.68 ± 0.56 | 0.35 ± 0.14 |
| V731CG1 | 12.22 ± 0.20 | 11.81 ± 1.14 | 37.90 ± 1.20 | -0.14 ± 0.26 |
| V731CG2 | 12.13 ± 0.18 | 12.41 ± 1.18 | 31.99 ± 0.94 | 0.36 ± 0.15 |
| V745CG1 | 5.20 ± 0.06 | 5.38 ± 0.38 | 32.53 ± 0.70 | 0.13 ± 0.10 |

**Table S5.** Methyl $S^2_{axis}\tau_c$ and $^1$H and $^{13}$C chemical shift anisotropies in FLN5, 283 K. $S^2_{axis}\tau_c$ values are tabulated from measurements of both $^{13}$C CCR (Fig. 3a) and $^1$H TQ build-up[1].

| Methyl | $\Delta\delta_H$ / ppm | $\Delta\delta_C$ / ppm |
|---|---|---|
| L661CD1 | 0.016 ± 0.009 | 0.39 ± 0.02 |
| L661CD2 | 0.027 ± 0.007 | 0.12 ± 0.02 |
| V664CG1 | 0.031 ± 0.006 | 0.36 ± 0.02 |
| V731CG1 | 0.005 ± 0.037 | 0.04 ± 0.05 |
| V731CG2 | -0.034 ± 0.007 | 0.41 ± 0.03 |
| I738CD1 | 0.039 ± 0.005 | 0.02 ± 0.02 |
| I743CD1 | -0.015 ± 0.004 | 0.48 ± 0.02 |
| V745CG1 | -0.027 ± 0.005 | 0.31 ± 0.02 |
| L701CD1 | -0.071 ± 0.028 | 1.09 ± 0.46 |
| V702CG1 | 0.059 ± 0.024 | 0.04 ± 0.03 |
| V702CG2 | 0.049 ± 0.021 | 0.54 ± 0.22 |
| V703CG1 | -0.081 ± 0.032 | 0.42 ± 0.17 |
| V703CG2 | 0.036 ± 0.016 | 0.27 ± 0.11 |

**Table S6.** Fitted chemical shift perturbations for FLN5 excited states, 283 K. Resonances are divided into two groups as indicated, with exchange parameters as shown in Fig. 6.

**Listing S1.** Bruker format pulse sequence for measurement of methyl Hahn echo DQ' and QQ relaxation.

```
/* Hahn echo relaxation measurement of four spin coherences in methyl groups
based on 1H TQ CPMG sequence (Yuwen, Vallurupalli & Kay, Angewandte Chemie, 2016)

    Relaxation times in vdlist
    Set td1 = 21 * number of relaxation times
    Apply receiver phase cycling post-acquisition

    Assumes that sample is specifically 13CH3 labeled

        1H: O1 on methyls (0.8 ppm)
            pwh = p1 1H pw90 @ power level pl1 highest power

       13C: O2 centre at 20 ppm
            pwc = p2 13C pw90 @ power level pl2 highest power
            power level pl21 is used for 13C decoupling.
*/

prosol relations=<triple>

#include <Avance.incl>
#include <Grad.incl>
#include <Delay.incl>

/************************/
/*   Define pulses     */
/************************/
define pulse dly_pg1     /* Messerle purge pulse */
  "dly_pg1=2m"
define pulse dly_pg2     /* Messerle purge pulse */
  "dly_pg2=3.4m"
define pulse pwh
  "pwh=p1"               /* 1H hard pulse at power level p1 (tpwr) */
define pulse pwc
  "pwc=p3"               /* 13C pulse at power level pl2 (dhpwr) */

/*************************/
/*   Define delays      */
/*************************/
"in0=inf2/2"
"d11=30m"

/*************************/
/*   Define f1180       */
/*************************/
  "d0=larger((in0)/2 - 2.0*pwh, 2e-7)"

define delay taua
  "taua=d3"              /* d3 ~ 1.8-2ms ~ 1.0s/(4*125.3)"  ~ 1 / 4J(CH) */
define delay taub
  "taub=d4"              /* d4 = 1/4JCH exactly */


"acqt0=0"              /* select 'DIGIMOD = baseopt' to execute */

aqseq 312

1 ze

2 d11 do:f2

  20u pl1:f1 pl2:f2


/******************/
/* Messerle purge */
/******************/
  20u pl11:f1
  (dly_pg1 ph26):f1
  20u
  (dly_pg2 ph27):f1
```

```
; off-resonance presat
30u fq=cnst10(bf hz):f1
30u pl9:f1
d1 cw:f1 ph26
4u do:f1
30u fq=0:f1
20u pl1:f1


/*******************************************/
/* Destroy 13C equilibrium magnetization */
/*******************************************/
  (pwc ph26):f2

  20u UNBLKGRAD

  2u
  p50:gp0
  d16

/************************/
/* Create QQ coherence */
/************************/

  (pwh ph1):f1

  2u
  p51:gp1
  d16

  "DELTA = taua - 2u - p51 - d16 - pwh*2.0/PI"
  DELTA

  (center (pwh*2 ph1):f1 (pwc*2 ph26):f2)

  2u
  p51:gp1
  d16

  "DELTA = taua - 2u - p51 - d16"
  DELTA

  (pwc ph3):f2

  2u
  p52:gp2
  d16

  "DELTA = taub - 2u - p52 - d16"
  DELTA

  (center (pwh*2 ph1):f1 (pwc*2 ph26):f2)

  2u
  p52:gp2
  d16

  "DELTA = taub - 2u - p52 - d16"
  DELTA

  (pwh ph1):f1

/*************/
/* Hahn echo */
/*************/
  vd*0.5
  (center (pwh ph29 pwh*2 ph26 pwh ph29):f1 (pwc*2 ph2):f2 )
  vd*0.5

/*****************************************************/
/* Begin back-transfer and chemical shift evolution */
/*****************************************************/
```

```
  (pwh ph26):f1

  2u
  p53:gp3
  d16

  "DELTA = taub - 2u - p53 - d16"
  DELTA

/********/
/* HMQC */
/********/
  (pwc*2 ph26):f2

  d0
  (pwh ph29 pwh*2 ph26 pwh ph29):f1
  d0

  2u
  p53:gp3
  d16

  "DELTA = taub - 2u - p53 - d16"
  DELTA

  (pwc ph4):f2

  "DELTA = pwc*2.0"
  DELTA

  (pwh ph27):f1

  2u
  p54:gp4
  d16

/*****************************************************************/
/* C->H back transfer, use wtg_flg for better water suppression */
/*****************************************************************/
  20u pl1:f1

  (pwh ph26):f1

  2u
  p57:gp7
  d16
  "DELTA = taua - 2u - p57 - d16 - p10 - 1u - larger(pwh,pwc) - pwh*2.0/PI"
  DELTA
  (p10:sp10 ph28):f1
  1u pl1:f1
  (center (pwh*2 ph26):f1 (pwc*2 ph27):f2 )
  1u
  (p10:sp10 ph28):f1
  "DELTA = taua - p57 - d16 - p10 - 1u - larger(pwh,pwc) - 2*pwc - 8u"
  DELTA
  p57:gp7
  d16

  4u BLKGRAD

  (pwc ph26):f2
  (pwc ph5):f2

  4u pl21:f2             /* lower power for 13C decoupling */

/*********************************/
/* Signal detection and looping */
/*********************************/
  go=2 ph31 cpds2:f2
  d11 do:f2 mc #0 to 2
    F1I(ip1, 7, ip3, 3)
    F1QF(ivd)
    F2PH(ip4, id0)
```

```
HaltAcqu, 1m
exit

ph0=1
ph1=(7) 0
ph2=0 2
ph3=(3) 0
ph4=0
ph5=0 2
ph26=0
ph27=1
ph28=2
ph29=3
ph31=0 2

;pl1 : tpwr - power level for pwh
;pl2 : dhpwr - power level for 13C pulse pwc (p2)
;pl9 : tsatpwr - power level for presat
;pl11 : tpwrmess - power level for Messerle purge
;pl21 : dpwr - power level for  13C decoupling cpd2
;p10 : 1000usec water flip-back
;sp10 : water flip-back (on H2O)
;spw14 : power level for eburp1 pulse
;spnam14: eburp1 pulse on water
;p1 : pwh
;p3 : pwc
;p14 : eburp1 pulse width, typically 7000u
;p50 : gradient pulse 50                                [1000 usec]
;p51 : gradient pulse 51                                [400 usec]
;p52 : gradient pulse 52                                [200 usec]
;p53 : gradient pulse 53                                [300 usec]
;p54 : gradient pulse 54                                [500 usec]
;p55 : gradient pulse 55                                [300 usec]
;p56 : gradient pulse 56                                [500 usec]
;p57 : gradient pulse 57                                [700 usec]
;pcpd2 : 13C pulse width for 13C decoupling
;d1 : Repetition delay D1
;d3 : taua ~1/(4*JCH)  ~1.8-2ms
;d4 : taub - set to 1/4JHC = 2.0 ms
;d11 : delay for disk i/o, 30ms
;d16 : gradient recovery delay, 200us
;cpd2 : 13C decoupling during t2 according to program defined by cpdprg2
;cpdprg2 : 13C decoupling during t2
;cnst10: water frequency for presat
;l1 : counter for the ncyc_cp values for cpmg
;l2 : actual value of ncyc_cp
;inf1 : 1/SW(X) = 2*DW(X)
;in0 : 1/(2*SW(x))=DW(X)
;nd0 : 2
;ns : 1*n
;FnMODE : States-TPPI, TPPI, States

;for z-only gradients:
;gpz0: 20%
;gpz1: 25%
;gpz2: 20%
;gpz3: -25%
;gpz4: 50%
;gpz5: -40%
;gpz6: -75%
;gpz7: -80%

;use gradient files:
;gpnam0: SMSQ10.32
;gpnam1: SMSQ10.32
;gpnam2: SMSQ10.32
;gpnam3: SMSQ10.32
;gpnam4: SMSQ10.32
;gpnam5: SMSQ10.32
;gpnam6: SMSQ10.32
;gpnam7: SMSQ10.32
```

**Listing S2.** nmrPipe and Julia processing scripts for analysis of DQ' and QQ Hahn echo experiments.

*proc.jl:*

```
#!/usr/bin/env julia

function proc(inputname, outputname, Δp1, Δp2)
    td = 2048
    nrelax = 14

    # input phase cycle
    ϕ1 = [0, 1, 2, 3, 4, 5, 6, 0, 1, 2, 3, 4, 5, 6, 0, 1, 2, 3, 4, 5, 6] * 2π / 7
    ϕ2 = [0, 0, 0, 0, 0, 0, 0, 1, 1, 1, 1, 1, 1, 1, 2, 2, 2, 2, 2, 2, 2] * 2π / 3
    nphase = 21

    # sum up echo and anti-echo pathways
    ϕrx1 = -Δp1*ϕ1 - Δp2*ϕ2
    ϕrx2 =  Δp1*ϕ1 + Δp2*ϕ2

    ϕrx1 = exp.(1im * ϕrx1)
    ϕrx2 = exp.(1im * ϕrx2)

    npoints = Int(filesize(inputname)/4 - 512)
    ncomplex = Int(npoints / (td*nphase*2))
    # preallocate data (and dummy header)
    header = zeros(Float32, 512)
    y = zeros(Float32, npoints)

    # read the input file
    open(inputname) do f
        read!(f, header)
        read!(f, y)
    end

    y = reshape(y, td, 2, :)
    yc = y[:,1,:] + 1im * y[:,2,:]

    y = reshape(yc, td, nphase, :)
    ϕrx1 = reshape(ϕrx1, 1, nphase, 1)
    ϕrx2 = reshape(ϕrx2, 1, nphase, 1)
    y1 = sum(y .* ϕrx1, dims=2)
    y2 = sum(y .* ϕrx2, dims=2)
    y = y1 + y2
    y = reshape(y, td, ncomplex)

    # read the file
    open(outputname, "w") do f
        write(f, header)
        for i=1:ncomplex
            write(f, Float32.(real.(y[:,i])))
            write(f, Float32.(imag.(y[:,i])))
        end
    end

    run(`sethdr $outputname -yN $nrelax -yT $nrelax`)
end

proc("cube.fid", "cubeQQ.fid", 3, -1)
proc("cube.fid", "cubeDQ.fid", 3, 1)
```

*nmrproc.com:*

```
#!/bin/csh

bruk2pipe -verb -in ./ser                                              \
  -bad 0.0 -ext -aswap -AMX -decim 1312 -dspfvs 21 -grpdly 76          \
  -xN               4096  -yN               294  -zN              128  \
  -xT               2048  -yT               294  -zT               64  \
  -xMODE             DQD  -yMODE           Real  -zMODE       Complex  \
  -xSW         15243.902  -ySW          294.000  -zSW        1912.046  \
  -xOBS          950.450  -yOBS           1.000  -zOBS        238.995  \
```

```
  -xCAR           0.400  -yCAR           0.000  -zCAR          16.700  \
  -xLAB              1H  -yLAB             Tau  -zLAB             13C  \
  -ndim               3  -aq2D         Complex                         \
-out cube.fid

# run Julia script to apply receiver phase cycling
./proc.jl

# relaxation times
set tauList = (0.1 1.0 2.0 3.0 5.0 7.0 10.0 13.0 16.0 22.0 29.0 37.0 46.0 56.0)

nmrPipe -in cubeQQ.fid -fn TP                          \
| nmrPipe  -fn ZTP                                     \
| nmrPipe  -fn TP                                      \
| nmrPipe  -fn SP -off 0.5 -end 1.00 -pow 2 -c 1.0    \
| nmrPipe  -fn ZF -auto                                \
| nmrPipe  -fn FT -auto                                \
| nmrPipe  -fn PS -p0 -83 -p1 0.00 -di -verb           \
| nmrPipe  -fn EXT -x1 1ppm -xn -0.7ppm -sw            \
| nmrPipe  -fn TP                                      \
| nmrPipe  -fn LP -fb                                  \
| nmrPipe  -fn SP -off 0.5 -end 1.00 -pow 2 -c 1.0    \
| nmrPipe  -fn ZF -zf 2                                \
| nmrPipe  -fn FT -alt -neg                            \
| nmrPipe  -fn PS -p0 58.00 -p1 180.00 -di -verb      \
| pipe2xyz -out ft/test%03d.ft2 -y -ov

sortPlanes.com -in ./ft/test%03d.ft2 -out ./ft/test%03d.ft2 -tau $tauList -title
xyz2pipe -in ft/test%03d.ft2 >cubeQQ.ft

nmrPipe -in cubeDQ.fid -fn TP                          \
| nmrPipe  -fn ZTP                                     \
| nmrPipe  -fn TP                                      \
| nmrPipe  -fn SP -off 0.5 -end 1.00 -pow 2 -c 1.0    \
| nmrPipe  -fn ZF -auto                                \
| nmrPipe  -fn FT -auto                                \
| nmrPipe  -fn PS -p0 -83 -p1 0.00 -di -verb           \
| nmrPipe  -fn EXT -x1 1ppm -xn -0.7ppm -sw            \
| nmrPipe  -fn TP                                      \
| nmrPipe  -fn LP -fb                                  \
| nmrPipe  -fn SP -off 0.5 -end 1.00 -pow 2 -c 1.0    \
| nmrPipe  -fn ZF -zf 2                                \
| nmrPipe  -fn FT -alt -neg                            \
| nmrPipe  -fn PS -p0 123.00 -p1 180.00 -di -verb     \
| pipe2xyz -out ft/test%03d.ft2 -y -ov

sortPlanes.com -in ./ft/test%03d.ft2 -out ./ft/test%03d.ft2 -tau $tauList -title
xyz2pipe -in ft/test%03d.ft2 >cubeDQ.ft
```

**Listing S3.** Bruker format pulse sequence for measurement of methyl [13]C CSA and $S_{axis}^2\tau_c$.

```
; 1H-coupled 13C HSQC with relaxation period for measurement of CCR
;
; with off-resonance presat
; ZZ/crusher periods, clean-up gradient pairs
; (90,-180) phase correction
; use baseopt
;

#include <Avance.incl>
#include <Delay.incl>
#include <Grad.incl>

"p2=p1*2"
"d2=p2"
"p4=p3*2"
"p22=p21*2"
"d4=1s/(cnst2*4)"
"d11=30m"
"d12=20u"
"d13=4u"

"in0=inf2"
"d0=4u"

"DELTA=d4-p16-d16-larger(p1,p3)-0.6366*p1"
"DELTA1=d4-p19-d16-p10-p1-4u-0.6366*p1"
"DELTA2=d4-p19-d16-p10-p1-12u"
"acqt0=0"

define delay vdMin
"vdMin = 2*p19 + 2*d16"

; calculate offset for WFB
"spoff1=cnst21-o1"

aqseq 312

1 ze
  vdMin
  d11 pl12:f2

2 d11 do:f2
  ; purge before d1
  20u pl6:f1
  (2mp ph1):f1
  (3mp ph2):f1

  ; off-resonance presat
  30u pl9:f1
  30u fq=cnst21(bf hz):f1
  d1 cw:f1 ph1
  30u do:f1
  30u fq=0:f1

  ; purge equilibrium 13C
  30u UNBLKGRAD
  4u pl1:f1 pl2:f2
  (p3 ph1):f2
  p16:gp0
  d16

  ; begin main sequence
  (p1 ph1)
  p16:gp1
  d16
  DELTA
  (center (p2 ph1) (p4 ph1):f2 )
  DELTA
  p16:gp1
  d16
  (p1 ph2)
```

```
  ; zz purge
  p16:gp2
  d16

  ; 13C t1
  (p3 ph11):f2
  d0
  "TAU = vd*0.5 - p19 - d16"
  TAU
  p19:gp5
  d16
  (p4 ph1):f2
  4u
  p19:gp5
  d16
  TAU
  (p3 ph12):f2

  ; zz purge
  p16:gp3
  d16

  ; final inept
  (p1 ph1)
  p19:gp4
  d16
  DELTA1
  (p10:sp1 ph3):f1
  4u pl1:f1
  (center (p2 ph1) (p4 ph1):f2 )
  4u
  (p10:sp1 ph3):f1
  DELTA2
  p19:gp4
  d16
  4u BLKGRAD
  4u pl12:f2

  go=2 ph31 cpd2:f2
  d11 do:f2 mc #0 to 2
    F1QF(ivd)
    F2PH(ip11, id0)

exit


ph1=0
ph2=1
ph3=2
ph11=0 2
ph12=0 0 2 2
ph31=0 2 2 0


;pl1 : f1 channel - power level for pulse (default)
;pl2 : f2 channel - power level for pulse (default)
;pl9 : f1 channel - power level for presaturation
;pl12: f2 channel - power level for CPD/BB decoupling
;p1 : f1 channel -  90 degree high power pulse
;p2 : f1 channel - 180 degree high power pulse
;p3 : f2 channel -  90 degree high power pulse
;p4 : f2 channel - 180 degree high power pulse
;p10 : f1 channel - 90 degree selective pulse [1000 usec]
;sp1 : f1 channel - 90 degree WFB (p10)
;d0 : incremented delay (2D)
;d1 : relaxation delay; 1-5 * T1
;d4 : 1/(4J)XH
;d11: delay for disk I/O                              [30 msec]
;d12: delay for power switching                       [20 usec]
;d13: short delay                                     [4 usec]
;cnst2: = J(XH)
;cnst21: off-resonance presaturation frequency (bf hz)
;inf1: 1/SW(X) = DW(X)
;in0: 1/SW(X) = DW(X)
```

```
;nd0: 1
;NS: 2 * n
;DS: 16
;td1: number of experiments
;FnMODE: States-TPPI, TPPI, States or QSEQ
;cpd2: decoupling according to sequence defined by cpdprg2
;pcpd2: f2 channel - 90 degree pulse for decoupling sequence

;for z-only gradients:
;gpz0: 46 %
;gpz1: 13 %
;gpz2: 17 %
;gpz3: 33 %
;gpz4: 29 %

;gradients
;p16: 1000u
;p19: 300u

;use gradient files:
;gpnam0: SMSQ10.100
;gpnam1: SMSQ10.100
;gpnam2: SMSQ10.100
;gpnam3: SMSQ10.100
;gpnam4: SINE.10
```

**Listing S4.** Bruker format pulse sequence for measurement of methyl $^1$H CSA.

```
;IPAP HMQC for measurement of 1H CSA via 1H CSA/1H-13C DD CCR
; set td1 = 2*number of relaxation time points

#include <Avance.incl>
#include <Grad.incl>
#include <Delay.incl>

"p2=p1*2"
"p4=p3*2"
"d2=1s/(cnst2*2)"
"d3=1s/(cnst2*8)"
"d11=30m"
"d12=20u"
"d13=4u"

"in0=inf2/2"
"d0=in0/2-0.63662*p3-2*p1"

; loop counter for IPAP
"l1=0"

define delay vdmin
"vdmin=2*(p2+4u+p17+d16)"

"acqt0=0"
baseopt_echo

aqseq 312

1 ze
  vdmin
  d11 pl1:f1 pl2:f2
2 d11

  20u
  "TAU1=vd*0.25-4u-p17-d16-p3"
  "TAU2=vd*0.25-p3-p1"
  "TAU3=vd*0.25-p1-4u-p17-d16-p3"
  "TAU4=vd*0.25-p3"

# ifdef OFFRES_PRESAT
    30u fq=cnst21(bf hz):f1
# endif /*OFFRES_PRESAT*/

  ; relaxation period
  d12 pl9:f1
  d1 cw:f1 ph29
  d13 do:f1
  d12 pl1:f1 pl2:f2
  30u fq=0:f1
  50u UNBLKGRAD

  (p3 ph1):f2    ; crush eq'm 13C magnetisation
  d13
  p16:gp1
  d16

  ; start main sequence
  (p1 ph1):f1  ; INEPT
  "DELTA1=d2-p16-d16+0.6366*p1"
  DELTA1
  p16:gp2
  d16

  ; purge element
  (p3 ph11):f2
  "DELTA=d3-p17-d16-larger(p1,p3)"
  DELTA
  p17:gp3
  d16
  (center (p2 ph1):f1 (p4 ph1):f2 )
  DELTA
```

```
  p17:gp3
  d16
  (p3 ph12):f2

  ; t1 evolution
  d0
  (p1 ph13):f1
  (p2 ph14):f1
  (p1 ph13):f1
  d0
  (p3 ph15):f2

  ; relaxation period
  "TAU=vd*0.5-p17-d16-p2-4u"
  TAU
  4u
  p17:gp4
  d16
  (p1 ph1):f1
  (p2 ph2):f1
  (p1 ph1):f1
  4u
  p17:gp4
  d16
  TAU

  ; IPAP back-transfer
  if "l1 % 2 == 0" {
    ; IP
    "DELTA2=d2*0.5-p16-d16-p3"
    p16:gp2
    d16
    DELTA2
    p4:f2 ph1
    "DELTA3=d2*0.5-p3-4u"
    DELTA3
    4u BLKGRAD
  } else {
    ; AP
    "DELTA2=d2-p16-d16-p3-4u"
    p16:gp2
    d16
    DELTA2
    p3:f2 ph1
    4u BLKGRAD
  }

  ; acquisition
  go=2 ph31
  d11 mc #0 to 2
    F1I(iu1, 2)
    F1QF(ivd)
    F2PH(ip15, id0)

  4u BLKGRAD
exit


ph1= 0
ph2= 1
ph11=0 2
ph12=1 1 1 1 3 3 3 3
ph13=0 0 1 1 2 2 3 3
ph14=1 1 2 2 3 3 0 0
ph15=0
ph29=0
ph31=0 2 2 0


;pl1 : f1 channel - power level for pulse (default)
;pl2 : f2 channel - power level for pulse (default)
;pl9 : f1 channel - power level for presaturation
;p1 : f1 channel -  90 degree high power pulse
;p2 : f1 channel - 180 degree high power pulse
```

```
;p3 : f2 channel -  90 degree high power pulse
;p4 : f2 channel - 180 degree high power pulse
;p16: homospoil/gradient pulse
;p17: gradient pulse [300 usec]
;d0 : incremented delay (2D)                       [3 usec]
;d1 : relaxation delay; 1-5 * T1
;d2 : 1/(2J)CH
;d3 : 1/(8J)CH
;d11: delay for disk I/O                           [30 msec]
;d12: delay for power switching                    [20 usec]
;d13: short delay                                  [4 usec]
;d16: delay for homospoil/gradient recovery
;cnst2: = J(CH)
;cnst21: frequency in Hz for off-res presat
;inf1: 1/SW(X) = 2 * DW(X)
;in0: 1/(2 * SW(X)) = DW(X)
;nd0: 2
;NS: 8 * n
;DS: 16
;td1: number of experiments
;FnMODE: States-TPPI, TPPI, States or QSEQ

;for z-only gradients:
;gpz1: 31%
;gpz2: 7%
;gpz3: -40%
;gpz4: 29%

;use gradient files:
;gpnam1: SMSQ10.100
;gpnam2: SMSQ10.100
;gpnam3: SMSQ10.32
;gpnam4: SMSQ10.32

;OFFRES_PRESAT: for off-resonance presaturation, set cnst21=o1(water)
;             option -DOFFRES_PRESAT (eda: ZGOPTNS)
```
